# Relaxation-selective Intravoxel Incoherent Motion Imaging of Microvascular Perfusion and Fluid Compartments in the Human Choroid Plexus

**DOI:** 10.1101/2024.12.09.626489

**Authors:** Chenyang Li, Zhe Sun, Jiangyang Zhang, Yulin Ge

## Abstract

The choroid plexus (ChP) plays an important role in the glymphatic system of the human brain as the primary source of the cerebrospinal fluid (CSF) production. Development of a non-invasive imaging technique is crucial for studying its function and age-related neurofluid dynamics. This study developed a relaxation-selective intravoxel incoherent motion (IVIM) technique to assess tissue and fluid compartments in the ChP in a prospective cross-sectional study involving 83 middle-aged to elderly participants (age: 61.5 ± 17.1 years old) and 15 young controls (age: 30.7 ± 2.9 years old). Using a 3T MRI scanner, IVIM, FLAIR-IVIM, LongTE-IVIM, and VASO-LongTE-IVIM were employed to measure diffusivity and volume fractions of fluid compartments and evaluate aging effects on microvascular perfusion and interstitial fluid (ISF). FLAIR-IVIM identified an additional ISF compartment with free-water-like diffusivity (2.4 ± 0.9 x10^-3^ mm^2^/s). Older adults exhibited increased ChP volume (2320 ± 812 mm^3^ vs 1470 ± 403 mm³, p=0.0017), reduced perfusion (6.5 ± 4.7 vs 3.6 ± 2.9 x10⁻³ mm²/s, p=0.0088), decreased ISF volume fraction (0.58 ± 0.09 vs 0.67 ± 0.06, p=0.0042), and lower tissue diffusivity (1.16 ± 0.14 vs 1.29 ± 0.17 x10^-3^ mm²/s, p=0.031) compared to younger adults. Relaxation-selective IVIM may offers enhanced specificity for characterizing age-related changes in ChP structure and fluid dynamics.

## 1. Introduction

Choroid plexus (ChP) is a highly vascularized structure situated within the ventricles of the brain (**Fig. 1a**). Its primary function is to produce the cerebrospinal fluid (CSF), which is essential to deliver nutrients, provide mechanical protection, and facilitate metabolic waste removal for the brain (1). Additionally, the ChP acts as the primary site of the blood-CSF barrier (BCSFB), playing a vital role in regulating the exchange of water between the capillary blood and CSF (2,3). Recent proteomic analyses of CSF samples have revealed that an enlarged ChP and its associated dysfunction could be a characteristic feature of certain subtypes of Alzheimer’s disease (AD) (4). This suggests that the ChP may have a significant role in the onset and progression of age-related neurodegenerative diseases (5,6).

**Figure 1.**
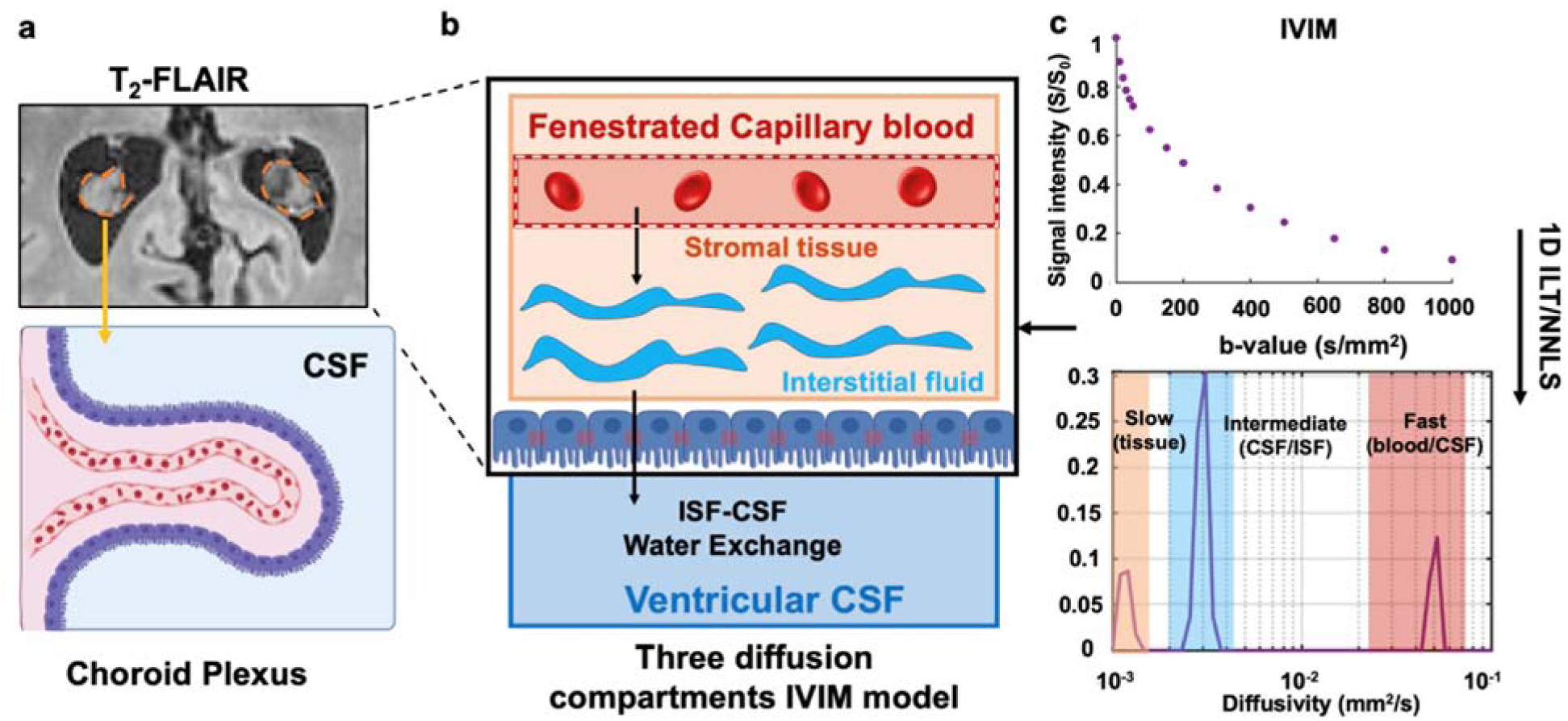
An overview of ChP functions and the three-compartment IVIM model used in this study. **(a)** The ChP is located in the ventricles and has a highly convoluted surface. **(b)** The production of CSF includes exchange between the vascular and stromal spaces and between stromal space and ventricles. **(c)** We hypothesize that the fluid compartments in the ChP can be studied using the three-compartment IVIM method enhanced with relaxation modulation.

The ChP produces CSF through a two-step process (**Fig. 1b**). Water, ions, and nutrients from the blood plasma first filter through the fenestrated capillaries into the stromal tissue, primarily driven by hydrostatic pressure. Following this process, the ChP’s epithelial cells selectively transport these elements into the ventricles, where they become CSF (2,3). To gain insights into CSF production, BCSFB functions, and waste clearance, it is key to assess the in vivo dynamics among blood, interstitial fluid, and CSF inside the ChP.

Currently, non-invasive evaluation of vascular functions of the ChP primarily uses contrast-based MRI techniques, including dynamic contrast enhanced MRI (DCE-MRI)(7,8), dynamic susceptibility contrast MRI (DSC-MRI)(9), and Ferumoxytol-enhanced susceptibility weighted imaging (SWI)(10). Contrast agents enhance the vascular space of the ChP, providing valuable information on vascular density, cerebral blood volume, and potentially BCSFB functions. However, gadolinium-based contrast agents can increase the risk of nephrogenic systemic fibrosis in elderly patients with kidney impairment. Therefore, it is preferable to use non-contrast techniques, such as arterial spin labeling (ASL) to assess vascular perfusion of ChP (11) and the functions of BCSFB (12–15). Due to ASL’s low resolution (∼2.5-3mm) and ChP’s small size, non-contrast assessment of structures and functions of the ChP, such as microvascular perfusion and other fluid dynamics (e.g. CSF or interstitial fluid (ISF)), remains challenging, as its special location within the CSF-filled ventricles lead to inevitable interference and substantial partial volume effects from the free ventricular CSF.

Intravoxel incoherent motion (IVIM) is a diffusion MRI technique sensitive to microvascular perfusion. Initially proposed by Le Bihan et al.(16–18), the IVIM model typically includes a fast diffusion component from blood flow in the capillary bed, known as pseudo-diffusion, a free water component in certain cases, and a slow diffusion component from tissue (**Fig. 1c**). IVIM has been extensively used to examine perfusion functions in the vascular-rich structures such as liver(19), kidney (20), and tumors(21). Given the ChP’s high vascularization and fenestrated capillary glomus, similar to the kidney, we hypothesize that IVIM can potentially be used to evaluate microvascular perfusion in the ChP. Furthermore, IVIM can also be extended for simultaneous evaluation of the ChP’s microstructural integrity, which is not available from contrast-enhanced MRI.

To date, only a limited number of studies have evaluated the feasibility of using IVIM to measure capillary perfusion(22) or assess the integrity of ChP tissue. In this research, we developed relaxation selective IVIM to simultaneously and separately analyze the tissue water, blood, and CSF compartments in the ChP, leveraging their distinct relaxation properties. Our techniques have demonstrated potential in identifying abnormalities and age-related changes in the ChP. Mapping of microvascular perfusion and other fluid compartments in the ChP could serve as an early diagnostic marker for assessing ChP functions in aging or age-related neurodegenerative diseases, such as Alzheimer’s disease and related dementias (AD/ADRD).

## 2. Materials and methods

### 2.1 Patient Characteristics

The study was approved by the Institutional Review Board. With HIPAA-compliant and IRB-approved acquisition policy, we recruited 15 young healthy controls (age: 30.7 ± 2.9, F/M=11/4), and 86 middle-aged to elderly subjects (age: 61.5 ± 17.1, F/M=52/31) recommended for MRI examination. All participants were given written, informed consent before MRI scans.

### 2.2 Pulse sequences

Clinical MRI examinations were performed on a Siemens 3.0T Prisma system using the 64-channel head coil. Clinical MR protocols include (a) 3D T_1_-magnetization prepared rapid gradient echo (T_1_-MPRAGE), 3D T_2_-fluid attenuated inversion recovery (T_2_-FLAIR) and susceptibility weighted images (SWI). T_1_-MPRAGE images were acquired with the following parameters: TE/TR = 2.96/2300ms; matrix size = 256×256×208; voxel size = 1×1×1mm^3^. T_2_-FLAIR images were acquired with TE/TR = 438ms/4800ms; matrix size = 256×256×160; voxel size = 1×1×1mm^3^. High resolution SWI was acquired with TE/TR = 22.5/38ms; matrix size = 600×768×36; voxel size = 0.31×0.31×1.5mm^3^.

IVIM experiments were performed using the standard diffusion-weighted echo planar imaging (EPI) sequence with matrix size of 192×192×20 and voxel size = 1.5×1.5×3 mm^3^. In this sequence, a 180-degree inversion pulse can be enabled before the diffusion encoding, with the inversion time (TI) modulating the T_1_-weighting (**Supplemental Fig 1-a**) in the acquired images. The echo time (TE) can also be adjusted to modulate the T_2_-weighting (**Supplemental Fig 1-b**). For example, the diffusion-weighted signal (*S*) in a homogeneous environment can be modulated by the TI and TE based on the following equation:

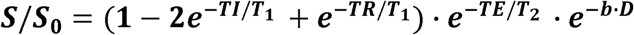

Where *S_0_*is the proton density, and *b* and *D* are the diffusion weighting and diffusivity, respectively.

### 2.3 Imaging the distributions of T_1_, T_2_ relaxation and diffusivity in the ChP

A series of T_1_-weighting EPI data, called mTI data here, were acquired using the sequence shown in **Supplemental Fig 1-a** with 19 inversion times (TI =0, 50, 325, 600, 875, 1150, 1430, 1800, 1980, 2250, 2530, 2800, 3080, 3350, 3630ms, 4180, 4450, 4730, and 5000 ms), TE/TR = 72/15000 ms, and no diffusion-weighting. A series of T_2_- weighted EPI data, called mTE data here, were acquired using the sequence shown in **Supplemental Fig 1-b** with 24 echo times (TE = 38, 43, 48, 53, 58, 63, 68, 73, 78, 83, 88, 93, 98, 103, 113, 123, 133, 143, 153, 163, 173, 183, 193, and 203 ms), TR = 8000 ms, and no diffusion weighting.

To obtain the distribution of diffusivity within the conventional IVIM framework, diffusion-weighted MRI data with fifteen b-values (0, 10, 20, 30, 40, 50, 100, 150, 200, 300, 400, 500, 650, 800 and 1000 s/mm^2^) were acquired at the same image resolution as the T_1_ and T_2_-weighted EPI data with TE/TR=58/8000ms. Representative mTI, mTE, and IVIM images are shown in **Supplemental Fig 2**, and the SNRs of the ChP were 371.4±60.1 (for the mTI images with TI=0), 397.9±124.2 (for the mTE images with TE=58ms) and 319.5±64.2 (for the IVIM images with b=0 s/mm^2^), as calculated by the mean of signal intensity in the ChP divided by standard deviation of background noise. The acquisition time of the mTI, mTE, and IVIM scans were 14, 15, and 2 minutes, respectively. The performance of the inversion pulse was also evaluated using a Siemens doped-water phantom with the same field of view and orientation as the human brain scans (**Supplemental Fig 3-a)**. The inversion efficiency (η) was fitted to evaluate the homogeneity of 180-degree inverted magnetization.

The mTI, mTE and IVIM signals were analyzed using inverse Laplace transform (ILT) using an open-source ILT package, which uses singular value decomposition with iterative optimization by Butler–Reed–Dawson methods(23). Regularization parameters of 10^-6^ were used to solve the distribution of diffusivities P(T_1_), P(T_2_) and P(D) within the range of TE and TI and b-value used and were calculated by integrating over all the peak amplitudes and further divided by the total peak amplitude to normalize the fraction from 0 to 1.

### 2.4 Conventional IVIM, FLAIR-IVIM, LongTE-IVIM and VASO-LongTE-IVIM

Given the distinct T_1_, T_2_ relaxation times of tissue, blood, and CSF, we proposed to selectively suppress one or two fluid compartments in order to enhance the sensitivity of IVIM to a particular fluid compartment. Specifically, we proposed three T_1_ and T_2_ selective IVIM acquisition schemes and compared it with the conventional IVIM acquisition.

1. Conventional IVIM scan were performed with fifteen b–values (0, 10, 20, 30, 40, 50, 100, 150, 200, 300, 400, 500, 650, 800, and 1000 s/mm^2^), diffusion direction along the slice direction, and TE/TR = 73/8000ms without the inversion preparation. The same selection of b-values and diffusion direction were used for relaxation selective IVIM.
2. Fluid attenuated inversion recovery (FLAIR)-IVIM were performed with an inversion recovery preparation targeting CSF with TI=1800 ms to suppress CSF signals, TE/TR = 73/8000ms.
3. LongTE-IVIM were performed with TE/TR = 183/8000ms and without the inversion preparation to suppress tissue signals, due to their relatively short T_2_ relaxation time, while maintaining blood and CSF signals.
4. VASO-longTE IVIM were performed with TE/TI/TR = 183/1150/8000ms. Inspired by vascular occupancy space fMRI (VASO-fMRI)(24,25), the selection of the TI to suppress blood was calculated given the previously reported blood T_1_ at 3T(26) (T_1,blood_ = 1624ms). Theoretically, VASO-longTE-IVIM only retains the CSF signal, which allows us to further explore the composition of fast diffusion components, assuming the blood signal is eliminated in this case. A single slice is used for this acquisition to avoid through plane flow artifacts in the ventricles.

Each sequence takes approximately 2 minutes. To account for T_1_ and T_2_ effects induced by IR and different TE, the diffusion data were normalized with their individual S_0_ values. To analyze the diffusion spectrum, 1D ILT was applied on all IVIM datasets to derive the diffusivity and its distribution P(D) of different subvoxel diffusion compartments. Among the 86 elderly subjects, all of which had FLAIR-IVIM scans, 25 subjects with additional LongTE-IVIM scans and 5 subjects for VASO-longTE-IVIM as a preliminary exploratory study. Representative images of the four IVIM acquisition schemes are shown in **Fig. 2**.

**Figure 2.**
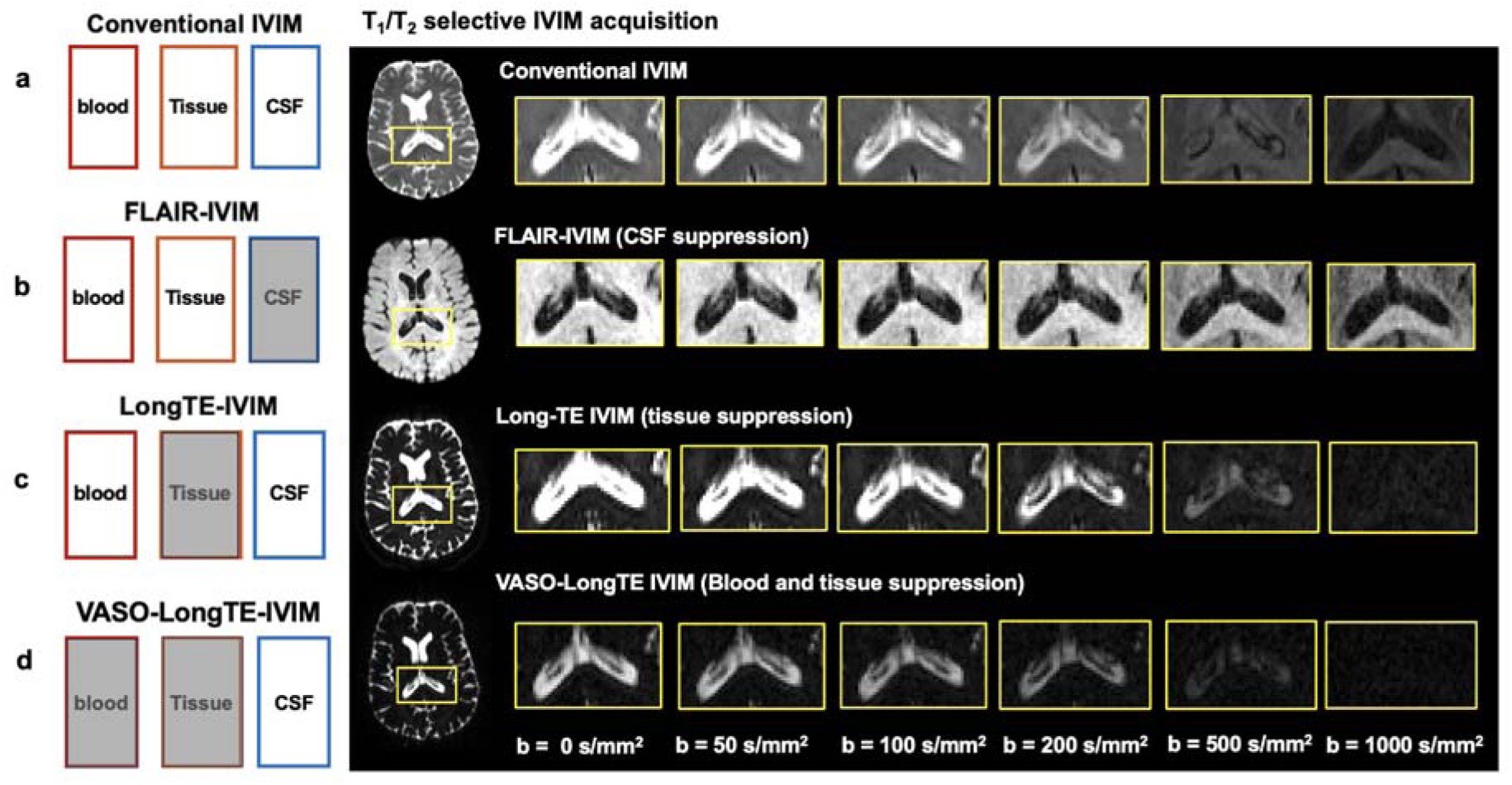
Illustration of T_1_ and T_2_ selective IVIM acquisition schemes used in this study and representative images showing the ChP. Conventional IVIM (**a**) includes signals from the blood, CSF, and tissue water compartments. FLAIR-IVIM (**b**) suppresses signals from the CSF compartments (shaded), as indicated by the dark ventricles in the non-diffusion-weighted (*b* = 0 s/mm^2^) image. LongTE IVIM (**c**) suppresses signals from the parenchymal tissue compartments, as indicated by the low tissue signals in the non-diffusion-weighted images. VASO-LongTE-IVIM (**d**) suppresses signals from both blood and parenchymal tissue compartments.

Following Wong et al.(27), a window from 1×10^-4^ to 1.5×10^-3^ mm^2^/s was used to calculate the fraction of slow diffusion in the parenchymal tissue. A window between 1.5×10^-3^ and 4×10^-3^ mm^2^/s was used to calculate the fraction of intermediate diffusion as it is within the region of free CSF or interstitial fluids, and a window above 4×10^-3^ mm^2^/s but lower than 1 mm^2^/s was used to calculate the fraction of the fast diffusion component (f_IVIM_).

### 2.5 Segmentation of the choroid plexus

The ChP was segmented from the T_1_-MPRAGE data using a Bayesian Gaussian mixture model (GMM) (https://github.com/EhsanTadayon/choroid-plexus-segmentation), which has shown improved segmentation accuracy to ChP(28). The ChP volume was calculated as the multiplication of voxel size and number of ChP voxel on T_1_-MPRAGE data.

### 2.6 Analytical Simulation

To better understand the signal evolution of different tissue types, we performed the analytical simulation to show the T_2_ signal decay and T_1_ signal recovery with respect to parenchymal tissue, CSF and blood to illustrate the fractional spin population changes and corresponding SNR under different acquisition schemes (**Supplemental Fig 4**). We analytically solved the T_1_ signal recovery by assuming the T_1,blood_=1560ms, T_1,CSF_=4000ms and T_1,tissue_=800ms. And T_2_ signal decay by assuming the T_2,blood_=250ms, T_2,CSF_=3000ms and T_2,tissue_=80ms. This simulation provided us with the approximate composition of spin populations and tissue specific SNR for different experiments.

## 3. Results

### 3.1 The tissue, blood, and CSF compartments in the ChP can be separated based on their distinct T_1_, T_2_, and diffusivity values

The T_1_ and T_2_ spectra from the ChP generated by ILT of the mTI and mTE data displayed three peaks with distinct T_1_ and T_2_ values. In the T_1_ spectrum (**Fig. 3a**), two peaks had T_1_ values (1794.0 ± 145.5ms and 4181.4 ± 579.7ms, respectively, n=10) that matched the T_1_s of blood and CSF in the literature (26). The other peak, with relatively shorter T_1_ values (249.4 ± 98.6ms, n=10), likely belonged to the ChP tissue. In the T_2_ spectrum (**Fig. 3c**), three peaks had T_2_ values corresponding to ChP tissue (31.8 ± 24.2ms), blood (194.7 ± 87.4ms), and CSF (3572.3 ± 1162.5ms). In comparison, T_1_ and T_2_ spectra from the lateral ventricles showed only one peak with T_1_ and T_2_ values of CSF (**Fig. 3b & d**).

**Figure 3.**
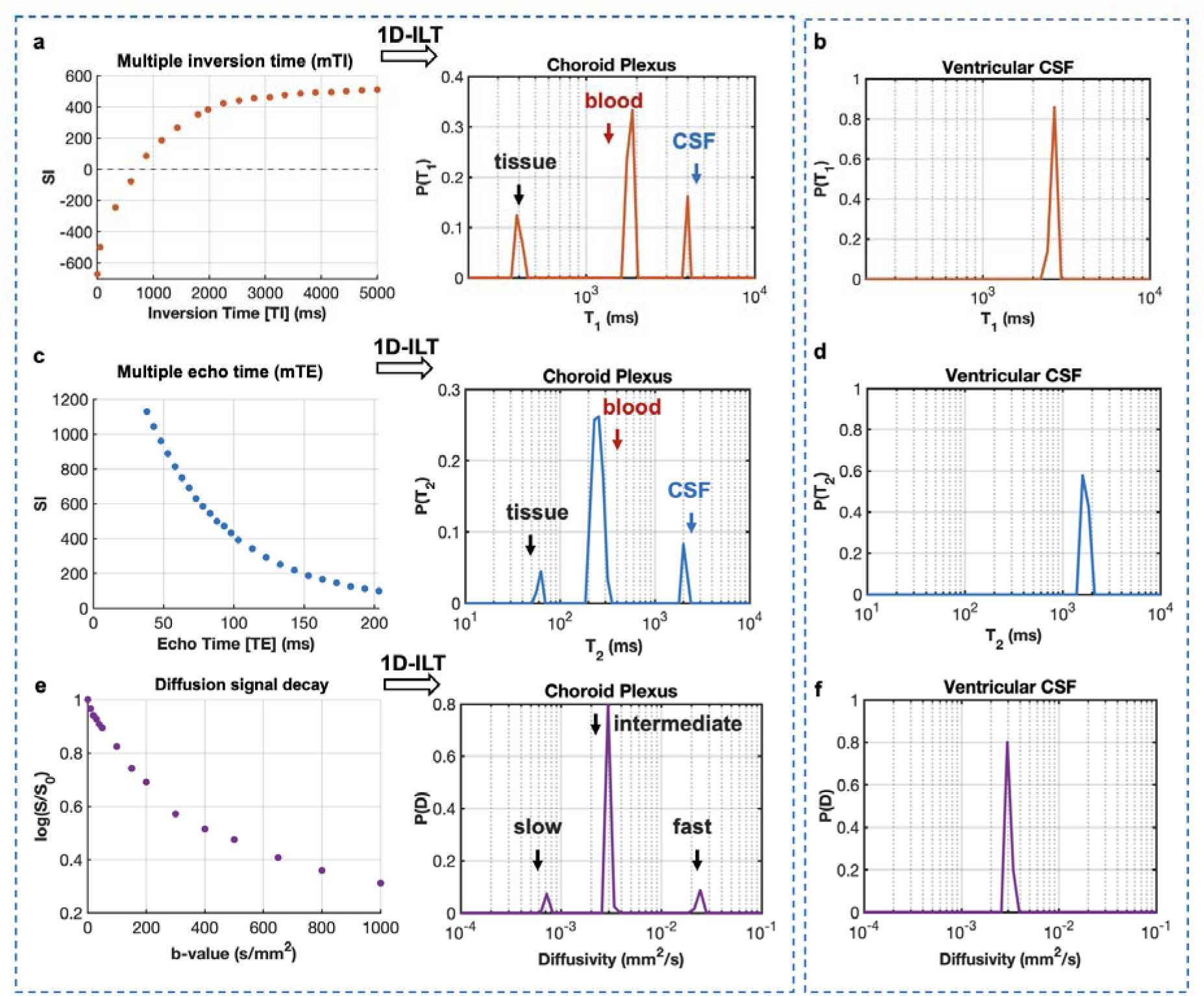
Representative multiple inversion time (**a**), multiple echo time (**c**), and IVIM (**e**) data from the ChP. ILT results show the three-peak distribution of T_1_, T_2_, and diffusivity in the ChP. In comparison, ILT results from the ventricular CSF (**b, d, & f**) only show one peak.

The IVIM data from the ChP also exhibited three distinct peaks after ILT (**Fig. 3e**). The peak with the fastest diffusivity (0.13 ± 0.11 mm^2^/s, n=10) was in the IVIM regime (flow component as the diffusivity is higher than free water). The peak with diffusivity close to those of free water (2.9±0.7×10^-3^ mm^2^/s, n=10) likely originated from ventricular CSF and water filled chambers within the ChP. The peak with the lowest diffusivity (0.7 ± 0.5×10^-3^ mm^2^/s, n=10) probably corresponded to the ChP tissue. In comparison, the IVIM data from the lateral ventricle (**Fig. 3f**) only showed one peak with diffusivity of free water.

Although mTE, mTI, and IVIM data all displayed three peaks in the ChP, it remained unclear whether there was a one-to-one relationship among the relaxivity and diffusion results. For example, in the T_2_ spectrum, the peak with T_2_ of the CSF had a lower volume fraction (0.17±0.09, **Table 1**) than the blood and tissue peaks (0.69±0.11 and 0.13±0.04, respectively, **Table 1**). Similar distribution was also observed in the T_1_ spectrum (**Table 1**). However, in the IVIM spectrum, the intermediate peak with diffusivity close to free water, presumably from CSF, had a higher volume fraction (0.85±0.13, **Table 1**) than the tissue and blood combined (0.096±0.03 and 0.05±0.02, respectively, **Table 1**). The difference suggested that the intermediate diffusion component included contributions from non-CSF sources.

**Table 1.** Distribution of T_1_, T_2_ relaxation time and diffusivity in the ChP with their corresponding volume fraction P(T_1_), P(T_2_) and P(*D*).

| Choroid Plexus | tissue | blood | CSF |
| --- | --- | --- | --- |
| $T_1$ (ms) | 249.4±98.6 | 1794.0±145.5 | 4181.4±579.7 |
| $P(T_1)$ | 0.13±0.04 | 0.69±0.11 | 0.16±0.08 |
| $T_2$ (ms) | 31.8±24.2 | 194.7±87.4 | 3572.3±1162.5 |
| $P(T_2)$ | 0.21±0.11 | 0.64±0.15 | 0.17±0.09 |

| Choroid Plexus | slow<br>(tissue water) | intermediate<br>(CSF/ISF) | fast<br>(flow) |
| --- | --- | --- | --- |
| $D(\times 10^{-3} \text{ mm}^2/\text{s})$ | $0.36 \pm 0.50$ | $2.9 \pm 0.7$ | $41.5 \pm 29.2$ |
| $P(D)$ | $0.096 \pm 0.03$ | $0.85 \pm 0.13$ | $0.05 \pm 0.02$ |

### 3.2 LongTE and FLAIR IVIM provide insights into the choroid plexus

To investigate the composition of the intermediate diffusion peak in the IVIM data, we first experimented with longTE acquisitions, in which signals from the short T_2_ peak in the T_2_ spectrum (31.8±24.2 ms, **Table 1**) should be mostly attenuated. The signal attenuation curve of longTE-IVIM showed a faster decay in the ChP than conventional IVIM (**Fig. 4a**). As expected, out of the three peaks in conventional IVIM (**Fig. 4d**), ILT of the LongTE-IVIM data from the ChP showed two peaks, without the slow diffusion peaks from conventional IVIM (**Fig. 4e**), suggesting the slow diffusion peak consisted of water with short T_2_, most likely in the tissue (**Fig. 4c**). As signals from tissue water were attenuated, the relative volume fractions of the fast and intermediate diffusion peaks changed from 0.05±0.02 to 0.49±0.17 and from 0.85±0.13 to 0.51±0.16, respectively, compared to conventional IVIM results (**Table 2**). The reduction in the relative volume fraction of the intermediate peak suggested that a portion of it might come from fluids with relatively short T_2_.

**Figure 4.**
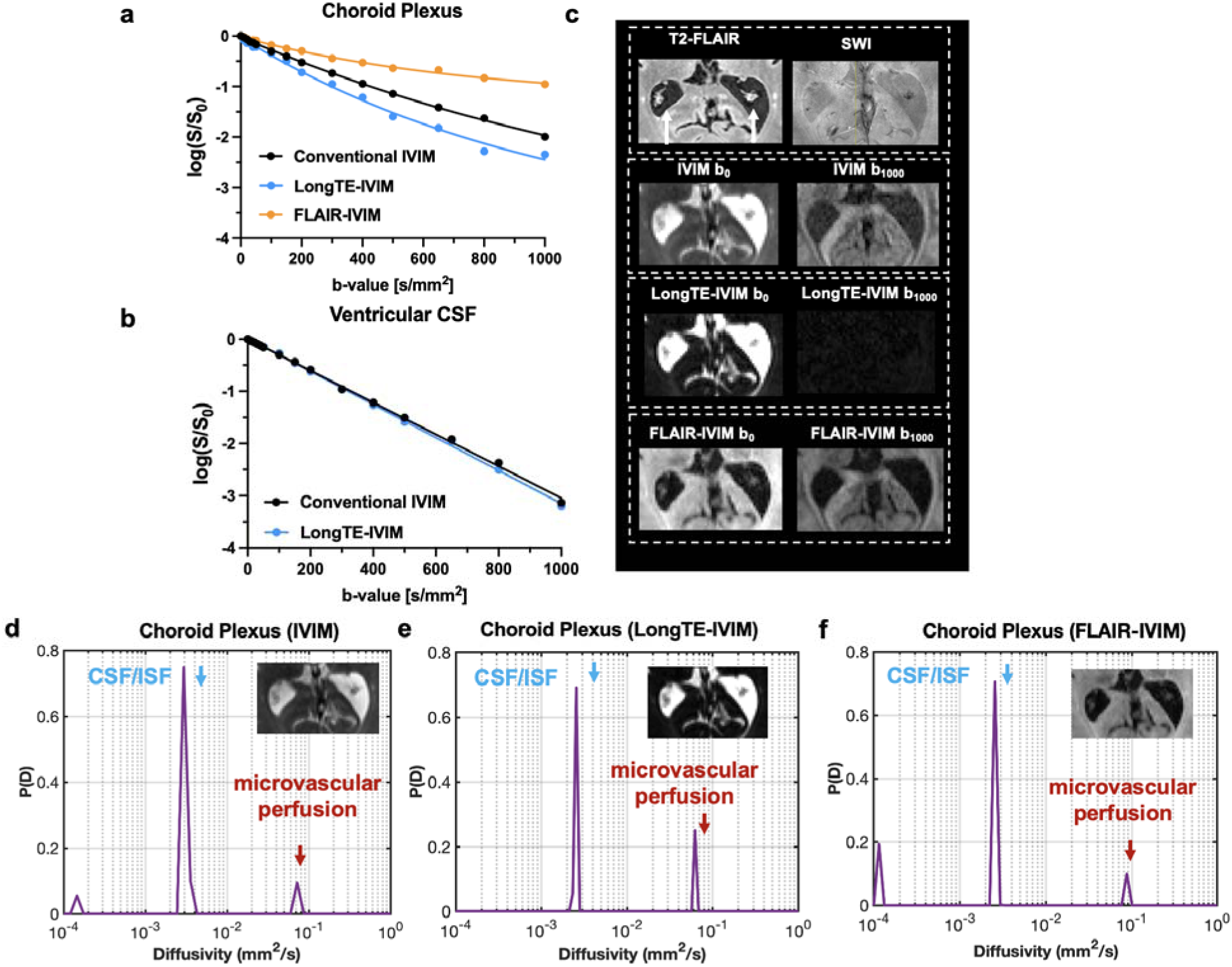
Representative log-scale IVIM signal decay in the ChP acquired using conventional IVIM, FLAIR-IVIM and LongTE-IVIM (**a**). In comparison, mono-exponential decay of CSF signal in the ventricles (**b**) with no apparent difference among results from the three IVIM method. (**c**) Representative images of T2-FLAIR, SWI and LongTE-IVIM b=0 and b=1000 s/mm^2^ images of IVIM, LongTE-IVIM, and FLAIR-IVIM. The spectral results showing the diffusivity distribution in the ChP from conventional IVIM (**d**), LongTE-IVIM (**e**) and FLAIR-IVIM (**f)**.

**Table 2.** Summary of diffusivity and its corresponding volume fraction in the slow, intermediate and fast diffusion components from FLAIR-IVIM and LongTE-IVIM.

| Choroid Plexus | $D_{\text{slow}}$<br>($\times 10^{-3} \text{ mm}^2/\text{s}$) | $D_{\text{int}}$<br>($\times 10^{-3} \text{ mm}^2/\text{s}$) | $D_{\text{fast}}$<br>($\times 10^{-3} \text{ mm}^2/\text{s}$) | $fD^*$<br>($\times 10^{-3} \text{ mm}^2/\text{s}$) |
| --- | --- | --- | --- | --- |
| FLAIR-IVIM | $0.28 \pm 0.31$ | $2.4 \pm 0.9$ | $54.1 \pm 36.8$ | $0.005 \pm 0.004$ |
| LongTE-IVIM | - | $2.8 \pm 1.3$ | $44.5 \pm 25.0$ | $0.024 \pm 0.019$ |

| Choroid Plexus | $f_{\text{slow}}$ | $f_{\text{int}}$ | $f_{\text{fast}}$ |
| --- | --- | --- | --- |
| FLAIR-IVIM | $0.27 \pm 0.13$ | $0.62 \pm 0.11$ | $0.09 \pm 0.05$ |
| LongTE-IVIM | - | $0.51 \pm 0.16$ | $0.49 \pm 0.17$ |

We then used the FLAIR preparation to suppress CSF and examine its impact to the intermediate peak. If the intermediate peak mainly came from free CSF, adding FLAIR will remove the intermediate diffusion peak. The signal attenuation curve of FLAIR-IVIM did show a slower decay than conventional IVIM (**Fig. 4a**). In the non-diffusion-weighted (b_0_) images, the ventricle appeared dark (**Fig. 4b**), suggesting near complete removal of signals from free CSF. Surprisingly, the ILT result from FLAIR-IVIM in the ChP still showed a substantial intermediate diffusion peak (**Fig. 4f**), with a volume fraction of 0.62±0.11 (**Table 2**), lower than in conventional IVIM but still higher than the slow and fast diffusion peaks combined (0.27±0.13 and 0.09±0.05, respectively, **Table 2**). These results suggested that the intermediate diffusion peak contained signals from both free CSF and fluid with T_1_ values distinct from CSF. In the ventricles, the signal attenuation curves acquired with the IVIM and LongTE-IVIM method showed no apparent difference (**Fig. 4b**). Signal attenuation in the ventricles of FLAIR-IVIM was not shown due to suppression of CSF signal. We did not combine FLAIR preparation with long TE IVIM due to poor signal to noise.

Apparent diffusion coefficient (ADC) value from the ChP tissue was calculated using the data with a b-value of 1000 s/mm^2^ and b_0_. The representative ADC map from IVIM and FLAIR-IVIM are shown in **Fig. 5a-b** and ADC map from IVIM and LongTE-IVIM are shown in **Fig.5c-d**. The FLAIR-IVIM showed a reduced tissue ADC of 1.2 ± 0.2 ×10^-3^ mm^2^/s compared to the conventional IVIM (2.5 ± 0.3 ×10^-3^ mm^2^/s, p<0.0001), whereas the longTE-IVIM data showed a higher tissue ADC value (2.8 ± 0.6 ×10^-3^ mm^2^/s). The difference was likely due to partial volume effects of the CSF, which was present in conventional and LongTE IVIM b_0_ images but suppressed in FLAIR-IVM b_0_ images.

**Figure 5.**
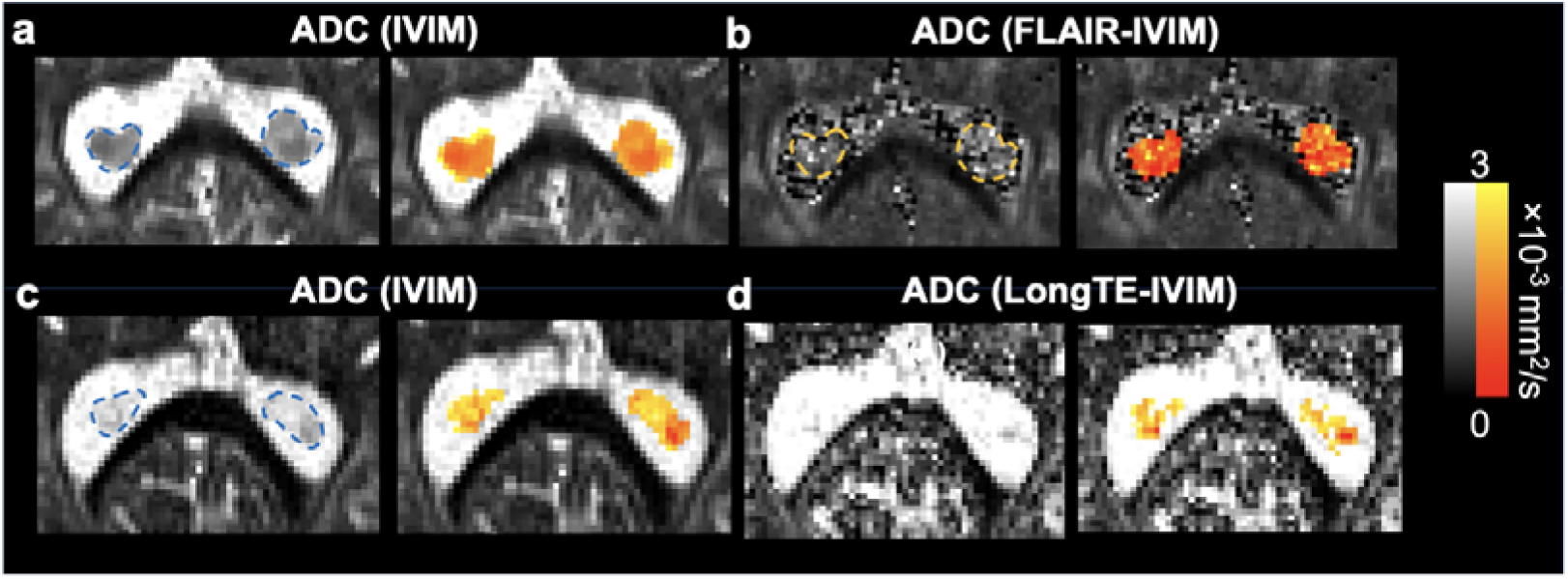
Representative ADC map estimated from conventional IVIM (a & c, different subjects), FLAIR-IVIM (b), and LongTE-IVIM (d).

### 3.3 VASO-LongTE-IVIM suggests the potential CSF/ISF flow in the ChP

We further explored the composition of the fast diffusion peak using the VASO- LongTE-IVIM, which should suppress most blood and tissue signals in the ChP (**Fig. 6a**). The data still revealed an intermediate peak (D = 3.2±0.6 ×10^-3^ mm^2^/s) with a volume fraction of 0.69 ± 0.23 (n=5) and a fast diffusion peak (D* = 34.5±29.3×10^-3^ mm^2^/s) with a volume fraction = 0.24 ± 0.18 (n=5) (**Fig. 6b**). The D* from VASO- LongTE-IVIM was lower than D* from conventional IVIM for the same subject. This remaining fast diffusion component with VASO preparation potentially indicates the presence of incoherent flow of other fluid with long T_2_, presumably CSF or ISF in the ChP (**Fig. 6e**).

**Figure 6.**
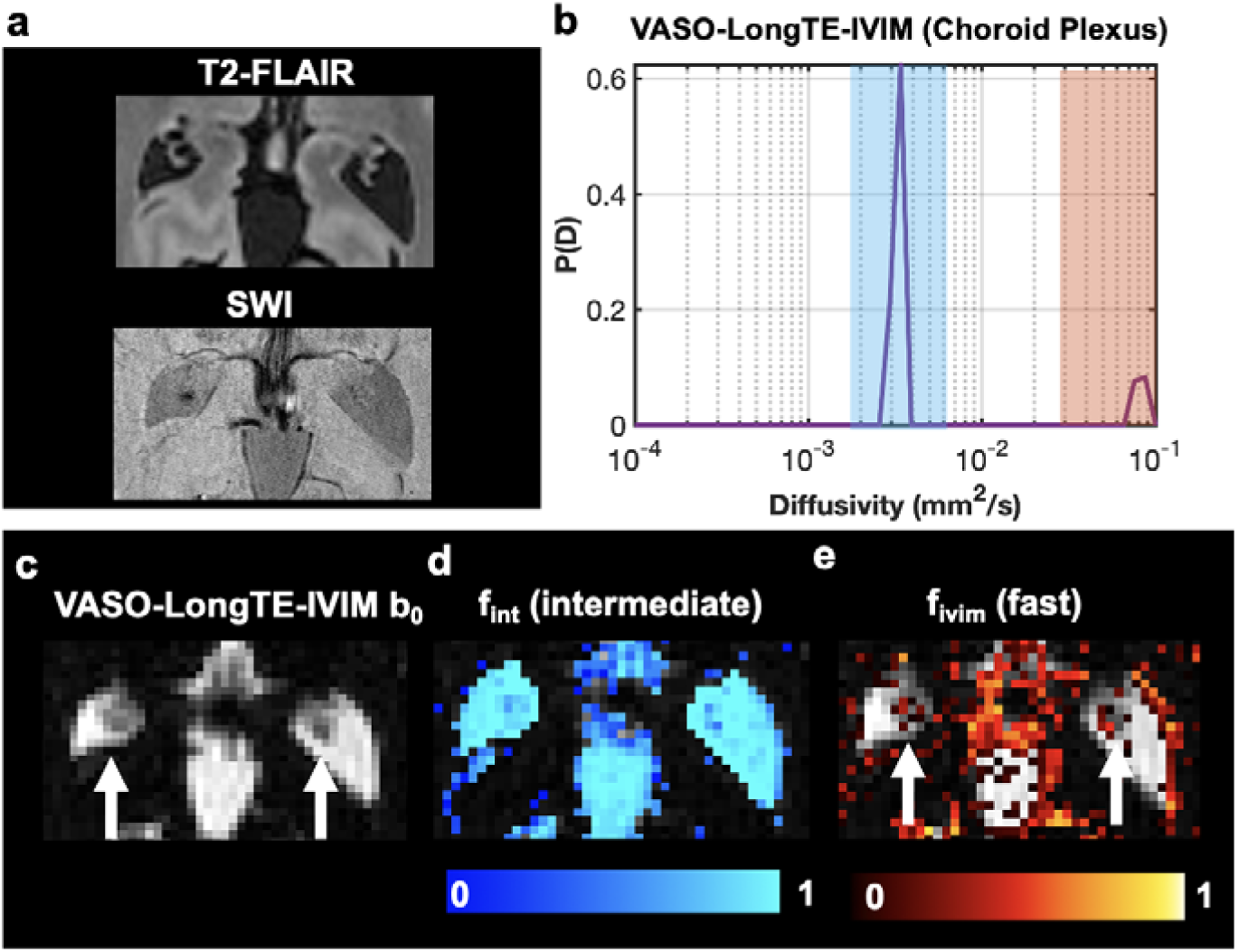
VASO-LongTE-IVIM of the ChP. (**a**) Representative T_2_-FLAIR and SWI for structural references of the ChP. (**b**) VASO-LongTE-IVIM spectral results of the ChP, with the intermediate diffusion peak (in the blue colored region) and fast diffusion peak (in the brass colored region). (**c**) VASO-LongTE-b0 images of the ChP. (**d**) voxel-wise mapping of the volume fraction of intermediate diffusion components. (**e**) voxel-wise mapping of volume fraction of fast diffusion components in the ChP.

### 3.4 Microvascular perfusion and the ISF compartment were reduced in elderly subjects

We used the FLAIR-IVIM technique to examine the ChP in elderly subjects. Defined in the 1mm isotropic resolution MPRAGE data, the volumes of the ChP in the elder subjects without apparent ChP abnormality (e.g. cyst) (N=46, age > 65 year old) were significantly higher than young subjects (N=15, age between 18- and 35-year-old) (**Fig. 7c**). There was no significant difference in diffusivity of the intermediate diffusion peak between the elderly and young subjects (**Fig. 7d**), but the volume fraction of the intermediate diffusion peak in the elderly subjects was significantly lower than the young subjects (**Fig. 7e**). For the fast diffusion peak, the elderly subjects showed significantly reduced *fD* and f_fast_* compared to the young subjects (**Fig. 7f-g**). Furthermore, the elderly subjects also showed significantly reduced tissue diffusivity compared to the young subjects (**Fig. 7h**).

**Figure 7.**
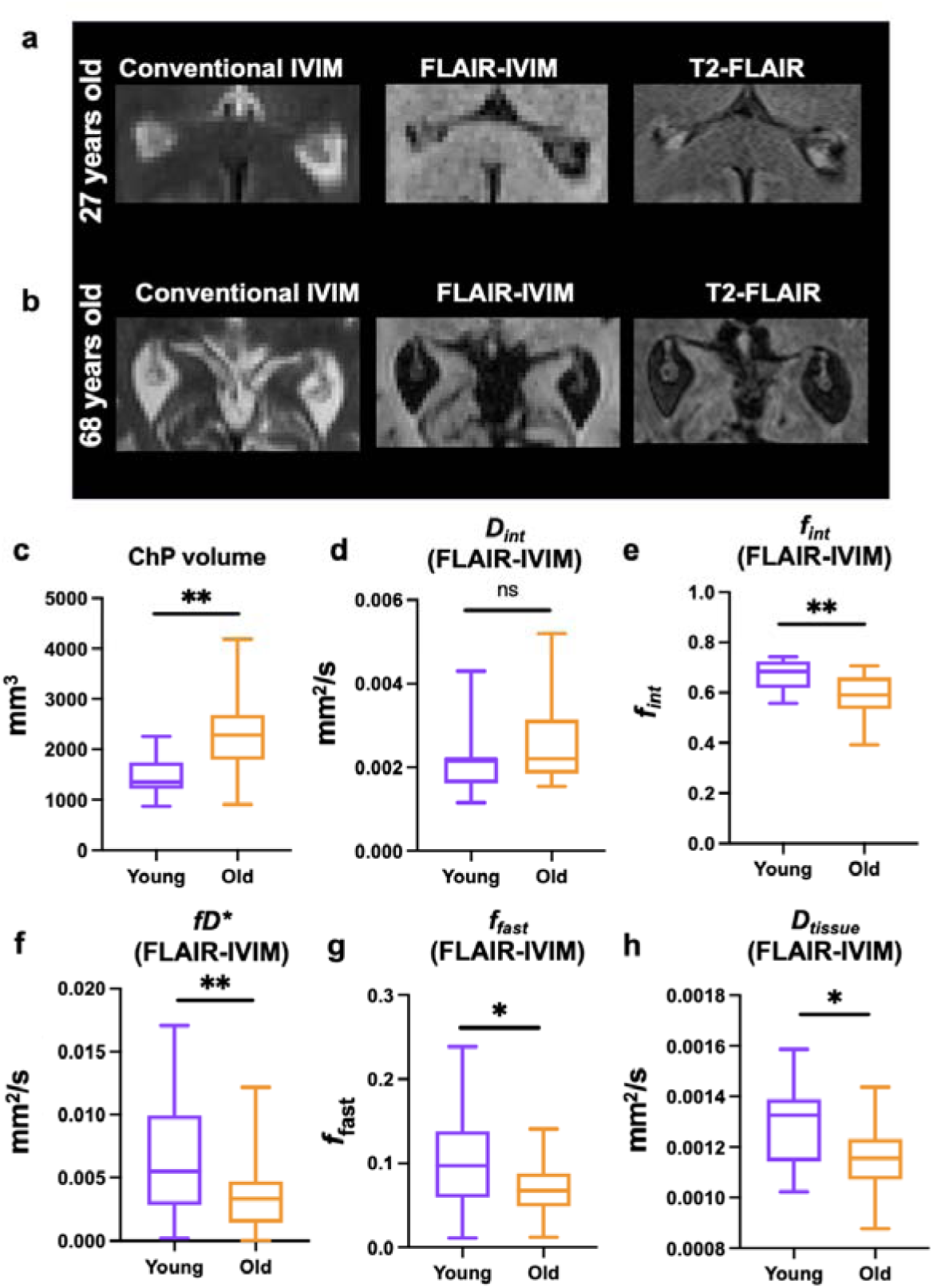
Comparisons of FLAIR-IVIM results from young and elderly subjects. (**a-b**) Representative IVIM, FLAIR-IVIM and T2-FLAIR images of a 27-year-old young adult subject (A) and a 68-year-old elderly subject (B). (**c**) Comparison of the ChP volume from T1-MPRAGE data between young (N=15) and elderly (N=46) subjects. Noted that the elderly population with apparent cyst structures were excluded for comparison. (**d-e**): Comparisons of the diffusivity and volume fraction of the intermediate diffusion peak between young and elderly subjects. (**f-g**) Comparisons of the fD*, volume fraction (f_fast_) of the fast diffusion peak between young and elderly subjects. (h) A comparison of tissue diffusivities between young and elderly. * and ** indicate significant difference between the two group with p < 0.05 and p < 0.01, respectively.

### 3.5 Choroid plexus cyst exhibit changes in relaxivity and diffusivity

Among 46 elderly populations, 11 elderly patients (age: 71.5±9.5, F/M =8/3) exhibited cysts or cyst-like structures in the ChP in T_1_-MPRAGE, T_2_-FLAIR, and SWI results. Among them, all showed reduced diffusivity, indicated by hyper-intense signals in diffusion-weighted images without FLAIR preparation (**Fig. 8**), compared to normal ChP (**Fig. 2**). Some cysts still showed unsuppressed signals in the T_2_-FLAIR images (**Fig. 8a**) as in normal ChP (**Fig. 4c**), whereas others were completely nulled in T_2_-FLAIR images (**Fig. 8b**), suggesting potential changes in relaxivities in the cysts. Under both cases the cyst-like structure was completely diminished in FLAIR-IVIM images at high b-values (e.g.,1000 s/mm^2^), suggesting the fluid compartment with reduced diffusivity had T_1_ values close to CSF.

**Figure 8.**
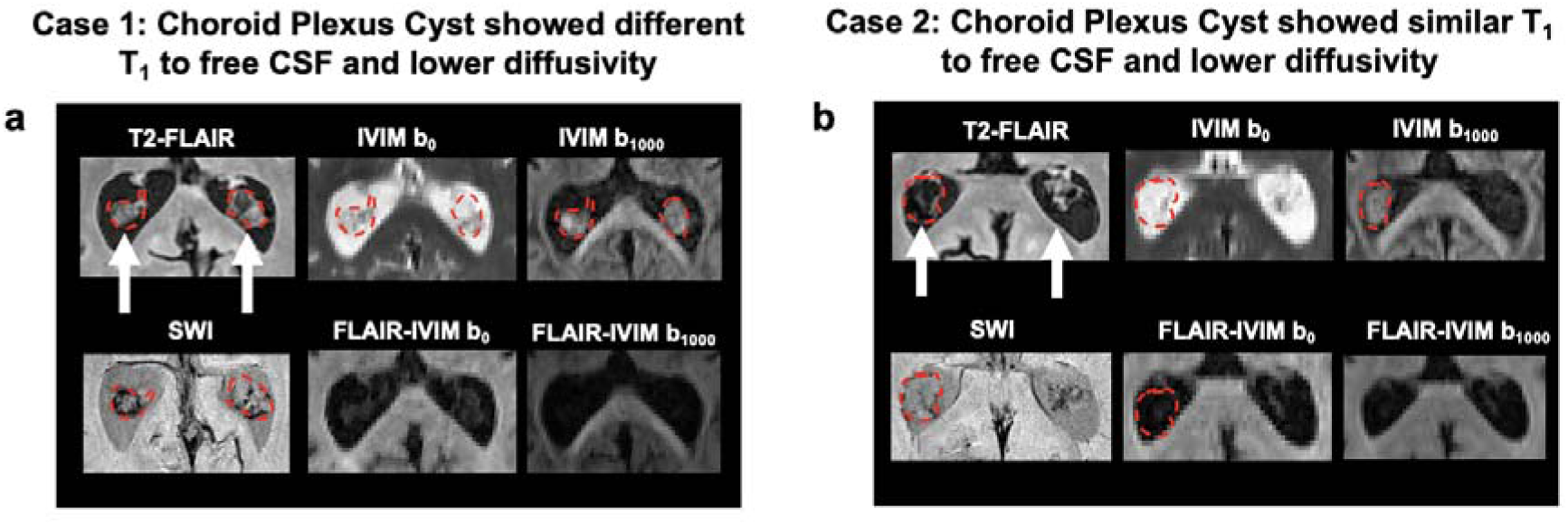
Representative cases of choroid plexus cysts in the elderly. In (**a**) case 1, the cysts exhibited restricted diffusion, and the cyst fluids was not fully suppressed by FLAIR and FLAIR-IVIM, while (**b**) the case 2 demonstrated that the cyst fluids were suppressed by FLAIR, indicating its T1 value was similar to ventricular CSF.

## 4. Discussion

The ChP, known for its critical role in CSF production, waste removal, and BCSFB function, has garnered increasingly attention, particularly regarding its link to age-related dementia. In this study, we introduced a relaxation-selective IVIM imaging technique designed to target the fluid compartments within the ChP based on their unique relaxivities. This method provides more specific information than conventional IVIM imaging. Our findings revealed a reduction in microvascular perfusion in the ChP of older subjects compared to young counterparts. Additionally, we discovered an ISF compartment within the ChP, which exhibits a reduced volume in older subjects. The relaxation-selective IVIM approach offers novel insights into the ChP’s substructure and its functional implications in CSF dynamics. With a rapid acquisition time of about 2 minutes of each of the IVIM acquisition scheme and a resolution of 1.5mm, these techniques are suitable to be combined with routine clinical scans and provide improved characterization of the ChP in neurodegenerative diseases.

### 4.1 Separating tissue, blood, and CSF in the choroid plexus

To determine the contributions of CSF, blood, and tissue water in each IVIM component, we utilized relaxation modulations, similar to spectral editing in NMR by selecting certain types of signals based on its relaxation properties. Previously, Jerome et al.(29) has proposed an extended T_2_-IVIM model to account for the TE dependence of the IVIM signal and demonstrated that the estimated perfusion fraction is dependent on the echo time used for IVIM. This finding highlights that relaxation effects are intrinsically encoded in the IVIM signal, and these effects can be modulated. To optimize IVIM acquisition, we tailored the TI and TE to detect specific fluid compartments. By selecting an appropriate TI, we were able to suppress CSF (FLAIR- IVIM)(30,31) and blood (VASO-LongTE-IVIM) signals (24), while a long TE helped suppressing tissue water (**Fig. 2**). This approach reduced the number of compartments under considerations and thereby allowed further examinations of the fluid compartments in the ChP. However, this approach only provides a partial view as IVIM reveals only relative volume fractions in diffusivity at given TI and TE, not their absolute volume fractions (**Fig. 3 & 4**), which are important for further quantification. Fully disentangling these components will may require T_1_-D and T_2_-D correlation data from advanced methods such as multi-dimensional MRI (MD-MRI)(32), which is time consuming due to the comprehensive sampling with multiple simultaneous relaxation and diffusion encodings and may not suit practical clinical use.

### 4.2 Evidence suggests an interstitial fluid compartment in the ChP

FLAIR-IVIM revealed a compartment in the ChP with diffusivities slightly lower than free water and T_1_ values different from CSF, likely representing ISF (**Fig. 4**). Wong et al. (27,33) previously observed similar intermediate diffusion peaks in the basal ganglia, cortex, and white matter hyperintensities (WMHs) in elderly patients with small vessel disease. They attributed these peaks to ISF, which may have different protein concentrations compared to CSF, causing the difference in T_1_. It was estimated that the human brain contains approximately 150 ml of ISF(2), with the ChP likely holding certain amount of ISF within its stromal tissue, derived from fenestrated blood vessels before being released into the ventricles as CSF. Anderson et al.(7) used a three water pools model for DCE-MRI to assess water exchange in the ChP between the blood and interstitial space and CSF release from interstitium into the ventricle. Our results suggested that the intermediate diffusion compartment observed in FLAIR-IVIM represented ISF in the ChP based on two observations: 1) the diffusivity of this fluid compartment was lower than those of free CSF and blood (**Table** 2); 2) the fluid compartment exhibited a shorter T_2_ than CSF and a different T_1_ from CSF, as indicated by results from LongTE-IVIM (**Fig. 4e**) and FLAIR-IVIM (**Fig. 4e**), suggesting a more restricted or viscous diffusion environment within the ChP tissue compared to the ventricles. Based on Einstein’s diffusion equation, the mean square distance of water molecule diffusion with the observed intermediate diffusivity and diffusion time of FLAIR- IVIM is approximately 20 µm, comparable to the size of cell bodies, suggesting the spatial scale of restrictive barriers within the stromal tissue. However, validating the ISF compartment through histopathology is challenging, as fluid is typically removed during conventional histological processing.

### 4.3 FLAIR-IVIM brings insights into age-related changes in the ChP

While the volume of the ChP generally increased with age (**Fig. 7c**), information on age-related changes in its internal structure and function is important to advance our knowledge on the effects of aging on ChP. Using FLAIR-IVIM, we reported reduced microvascular flow in elderly subjects (**Fig. 7f-h**). This reduction in vascular density in aging choroid plexus has been observed in high-resolution Ferumoxytol-enhanced SWI(10).

Furthermore, we observed a reduced intermediate diffusion fraction in the ChP in elderly subjects compared to young adults (**Fig. 7e**), which could indicate either actual volume changes of the ISF or alterations in protein or plasma concentration that affect the T_1_ of the ISF. Nevertheless, it has been reported that the CSF production rate is reduced with age due to several structural alterations in the ChP. With increasing age, the capillary wall tends to get thicker, affecting the exchange between capillary blood water and interstitial fluid, as a result, the composition of ISF in ChP may be altered. Additionally, we observed reduced tissue ADC using FLAIR-IVIM, the interstitial space of stromal tissue may undergo increased stromal sclerosis, calcification, and cyst formation. These changes could lead to a decrease in both tissue water diffusivity and fluid volume within the interstitial space of the ChP, thereby affecting CSF production efficiency in the elderly. However, other than diffusion, the interpretation of these results may be affected by the age-related change in tissue or interstitial fluid relaxivity, as remaining spin fractions of tissue signal could also affect the normalized volume fraction of interstitial fluids.

FLAIR-IVIM also helped to understand the cyst or cyst-like structures in the ChP. While conventional and FLAIR-IVIM results showed reduced diffusivities in the cysts compared to normal ChP, FLAIR-IVIM further suggested altered relaxivity in the cysts (**Fig. 8**) hinting a more restrictive physical environment and further changes in the chemical environment, respectively. Using oscillating gradient spin echo (OGSE)- diffusion MRI, Maekawa et al.(34) have found that ChP cysts demonstrated restrictive diffusion at shorter diffusion times, indicating that cyst fluids is spatially restricted or highly viscous due to higher protein levels(35,36). These observations have demonstrated the complexity of physical and physiochemical environment of the ChP cyst.

### 4.4 Evidence of potential CSF/ISF flow in the ChP from VASO-LongTE-IVIM

Given the CSF production process in the ChP, IVIM could potentially detect CSF or ISF movement within the ChP. Recent reports have shown that diffusion MRI with low diffusion-weightings may be sensitive to CSF flow. For example, Harrison et al. (37) reported the use of long echo times and low diffusion-weightings to detect perivascular fluid flow near the rat middle cerebral artery. Wen et al. (38,39) used a low b-value of 150 s/mm^2^ to measure slower CSF flow with suppression of fast nearby blood flow. Because the surface of ChP is highly folded, the CSF/ISF flow may appear incoherent and fall in the IVIM regime, which may provide complementary insights to ChP functions. VASO-LongTE-IVIM aims to null the signal from both blood and tissue components before IVIM readout, with the residual pseudo-diffusion component theoretically coming from the CSF/ISF flow (**Fig. 6**). However, these findings need further validations. One alternative explanation is that the VASO inversion pulses are calculated based on agreed a single T_1,blood_ at 3T, making it difficult to confirm that the blood signal is 100% suppressed. Therefore, the fast diffusion components we see may still contain signals from unsuppressed blood.

### 4.5 Technical considerations and limitations

For FLAIR-IVIM and VASO-IVIM, an important factor that may affect the image acquisition is the B_1_ inhomogeneities. Our water phantom result showed that the inversion efficiency is between 97 and 100% across the field of view. In our study, we assumed uniform inversion efficiency in the ChP to estimate T_1_ distributions. However, as noted by Avaram et al.(32), other dynamic processes, such as magnetization transfer, can impact short T_1_ components in inversion recovery experiments (40). Their impact on ChP tissue needs further investigation. In addition, assuming isotropic microstructural organization in the ChP, our FLAIR-IVIM protocol had only one diffusion direction, additional diffusion encoding directions will likely improve IVIM results but prolong acquisition times. In the lateral ventricles, flow void artifacts increase with b-values. Using flow-compensated techniques in the future could help mitigate these artifacts. Moreover, this study did not account for the exchange between different fluid compartments. Since water exchange across tissue compartments is common and crucial for the BCSFB function, future IVIM experiments with varying diffusion times could provide better insights into fluid compartment interactions in the ChP.

## 5. Conclusion

In summary, we developed relaxation-selective IVIM acquisitions to assess the structure and function of the ChP by targeting specific fluid compartments. Beyond the CSF, blood, and tissue compartments identified with conventional IVIM, FLAIR-IVIM results revealed an additional ISF compartment within the ChP. In elderly subjects, relaxation-selective IVIM demonstrated reduced microvascular flow, a diminished ISF compartment, and lower tissue ADC values compared to younger adults. This technique has the potential to elucidate age-related changes in ChP function and neurofluid dynamics in the brain.

## Notes

### Competing Interest Statement

The authors have declared no competing interest.

