## Supplemental Figures for "Relaxation-selective Intravoxel Incoherent Motion Imaging of Microvascular Perfusion and Fluid Compartments in the Human Choroid Plexus"

**Supplementary materials**:

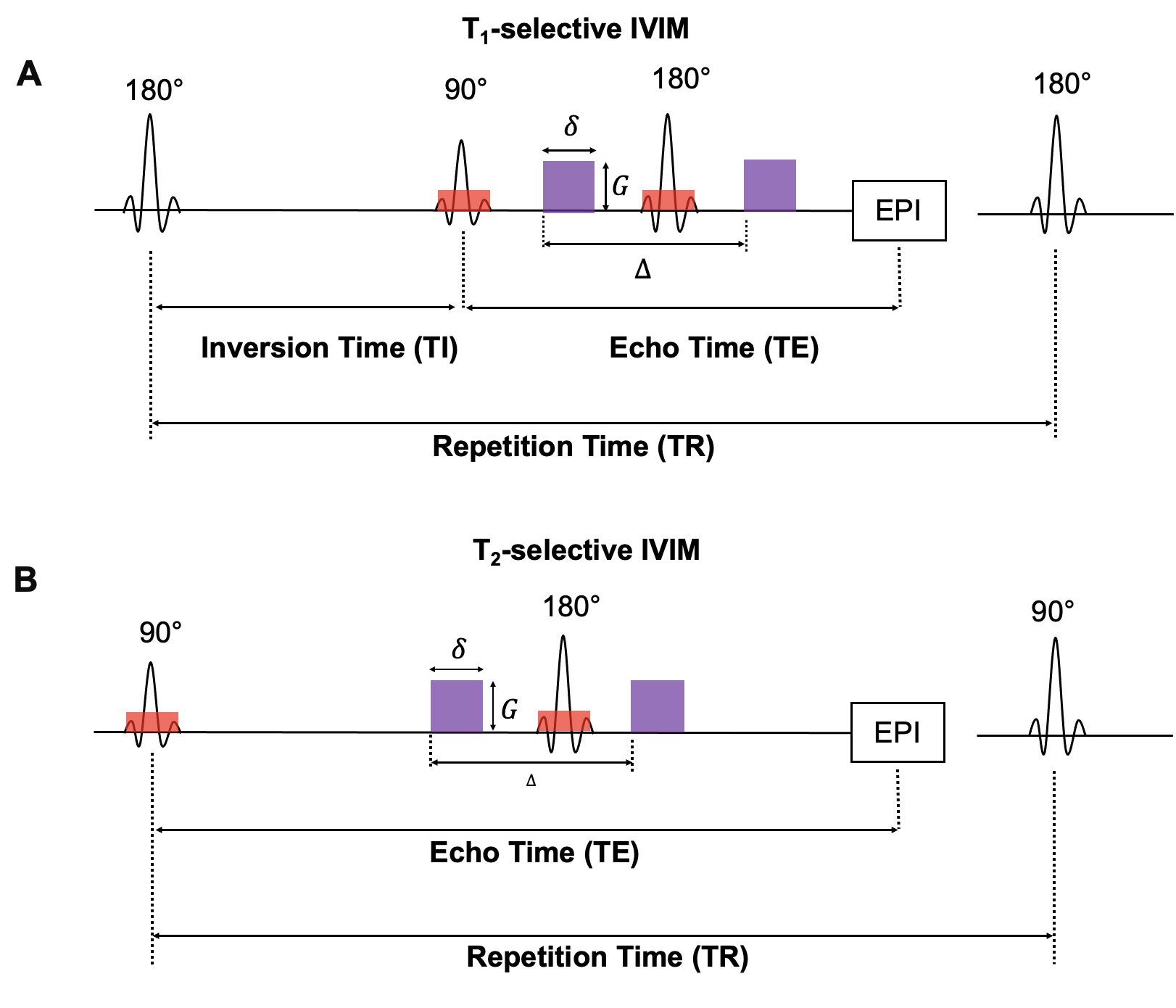

**Supplemental Figure 1.** Pulse sequences of T_1_ and T_2_ relaxation selective diffusion MRI acquisition. (**a**) 180 inversion pulse were applied before the IVIM encoding, adjusted inversion time (TI) was applied for T_1_-selective IVIM. (**b**) T_2_ selective IVIM was achieved by adjusting the echo time (TE) for T_2_-weighting.

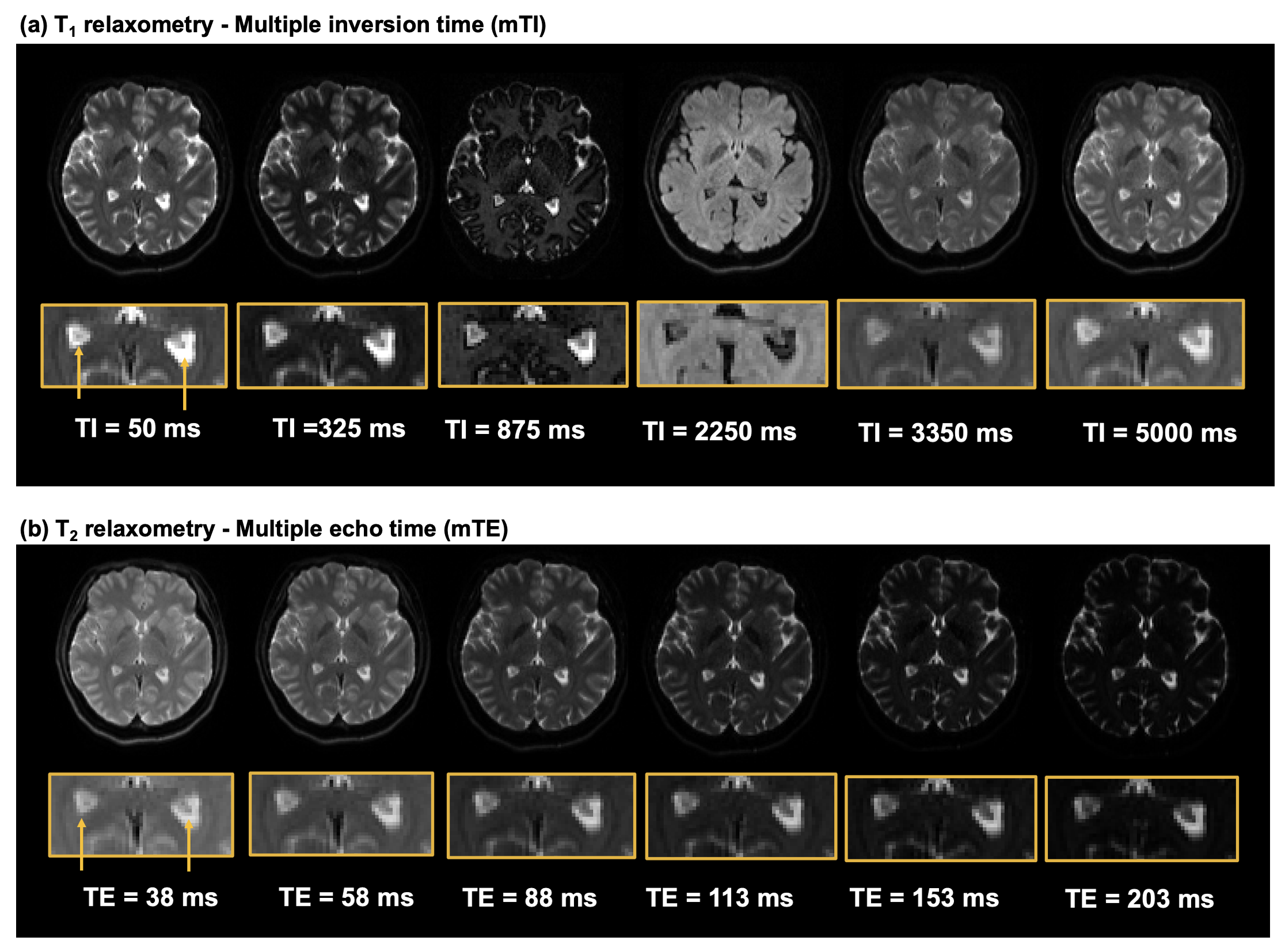

**Supplemental Figure 2**. Representative images of (a) multiple echo time for T_2_ relaxation distribution and (b) multiple inversion time for T_1_ relaxation distribution.

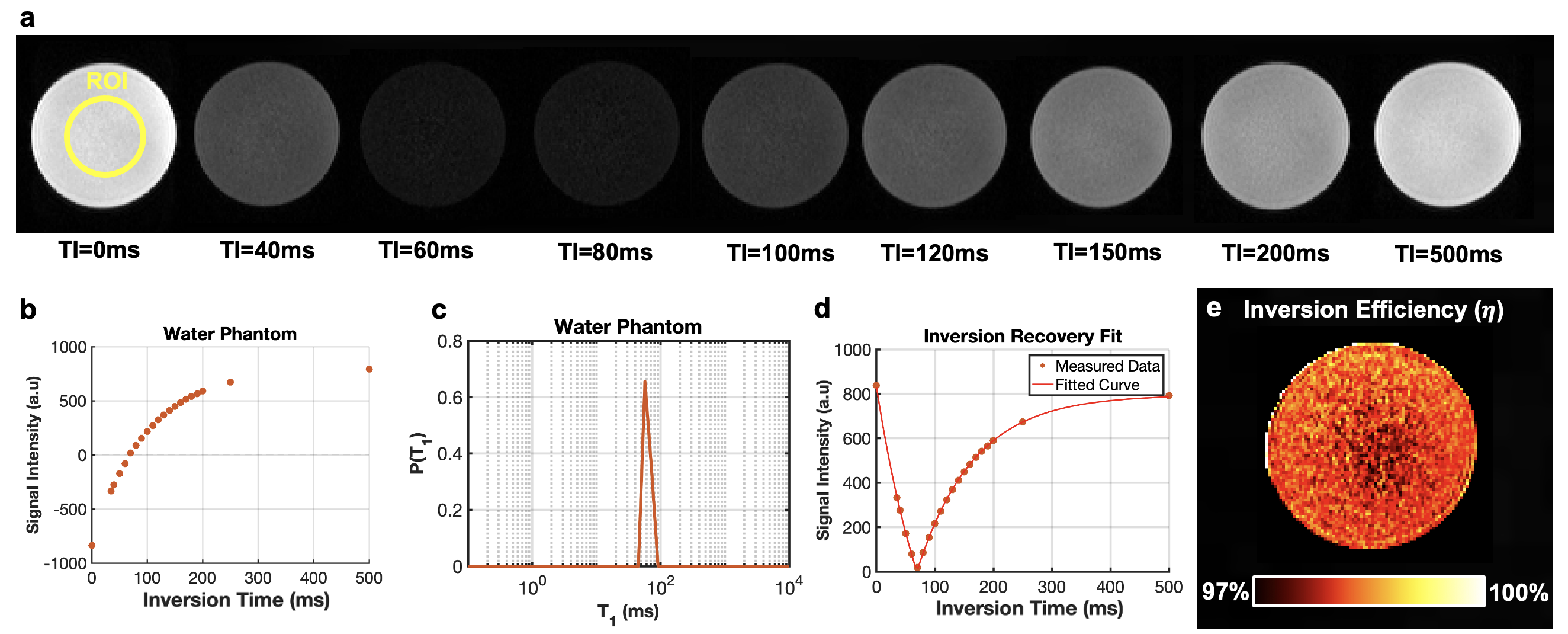

**Supplemental Figure 3**. (**a**) Multiple inversion time on a doped-water phantom from 0ms to 500ms due to short T_1_ of doped water compared to pure water. (**b**) Phase-corrected inversion recovery data prepared for ILT analysis. (**c**) ILT result of water phantom revealed doped water T_1_ of 91ms. (**d**) fitted T1 inversion recovery data and (**e**) voxel wise inversion efficiency map ($\eta$).

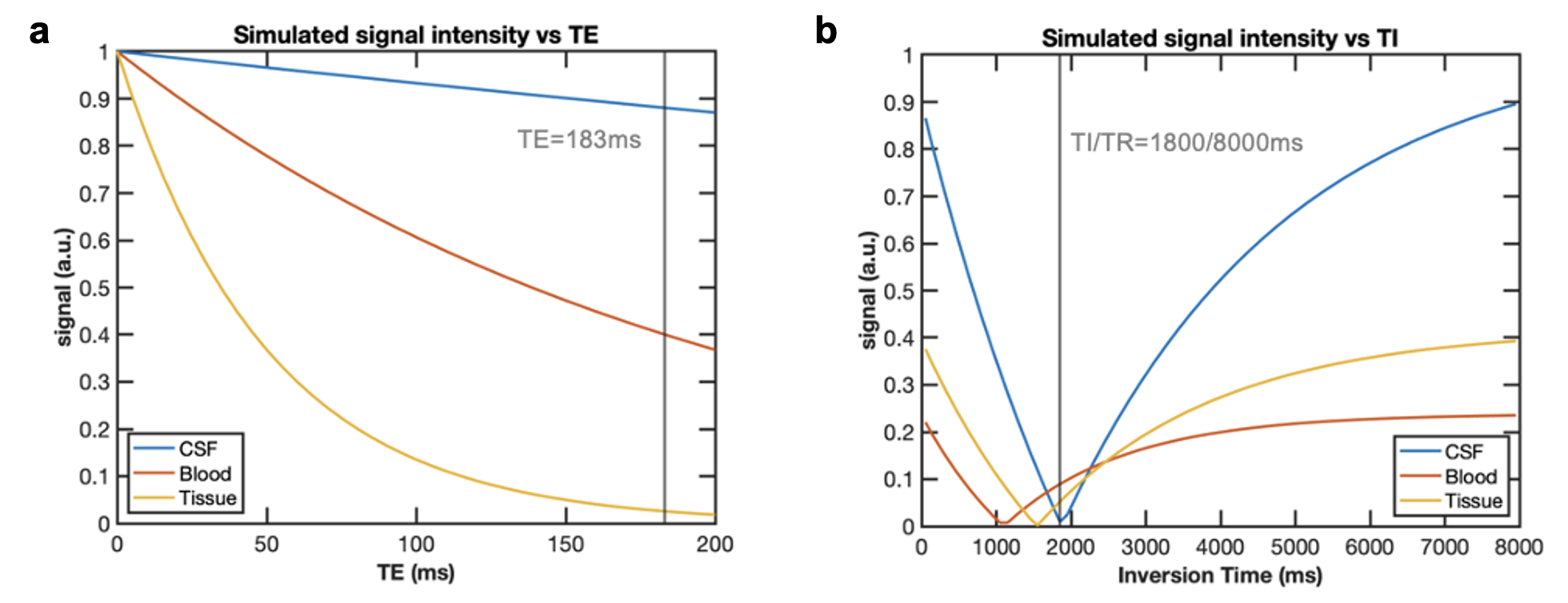

**Supplemental Figure 4**. Analytical simulation on (**a**) T_2_ signal decay and (**b**) T_1_ signal recovery in parenchymal tissue, CSF and blood showing spin populations at certain inversion times and echo times.
